# *Bacillus subtilis* YpeB holds SleB inactive preventing cortex peptidoglycan degradation during spore dormancy

**DOI:** 10.64898/2026.06.01.729290

**Authors:** Yongqiang Gao, Jeremy D. Amon, Joshua C. Cofsky, David Z. Rudner

## Abstract

Bacterial endospores are encased in a thick layer of specialized peptidoglycan called the cortex that is essential for core dehydration and heat resistance. Spore germination and outgrowth requires cortex degradation by enzymes that are deposited in the spore during sporulation. How these enzymes are held inactive during dormancy and activated during germination remains poorly understood. In *Bacillus subtilis* and many other endospore-forming bacteria, one of the lytic enzymes, SleB, is encoded in an operon with its putative regulator YpeB. The two proteins depend on each other for stability, but whether and how YpeB inhibits SleB were unknown. AlphaFold predicts a high-confidence interaction between the two proteins with YpeB’s PepSY domains embracing SleB’s catalytic domain. Here, we demonstrate that the two proteins can be co-purified when expressed in *E. coli*. Furthermore, in *B. subtilis* spores, amino acid substitutions at the predicted SleB/YpeB interface destabilized both proteins, resulting in impaired spore germination in the absence of the functionally redundant cortex lytic enzyme CwlJ. Selection for germination-competent suppressors identified general and allele-specific suppressors in *sleB* or, separately, *ypeB* that stabilized both proteins. Altogether, our data support a model in which YpeB inhibits SleB in the dormant spore through direct interaction. We propose that YpeB’s PepSY domains function as chaperone-inhibitors of SleB, akin to the role of pro-domains in protease zymogens. Chaperone dependence ensures that SleB proteins that fail to interact with YpeB remain unfolded and are ultimately degraded, while YpeB-bound SleB enzymes persist but are inhibited, preventing inappropriate cortex degradation during dormancy.

**IMPORTANCE:** Bacterial spores are among the most resilient cell types in nature. Their ability to resist sterilization during dormancy yet rapidly germinate and resume growth are central to the transmission and pathogenesis of spore-forming pathogens. A key component of their resistance is a thick layer of specialized peptidoglycan called the cortex that encases them. The cortex maintains the spore core in a highly desiccated state by physically restricting its expansion. Germination and outgrowth requires the degradation of this essential envelope layer. The enzymes responsible for cortex degradation are deposited in the spore during sporulation. How these lytic enzymes are held in active during dormancy has been a longstanding question. One of these enzymes, SleB, is encoded in the same operon as its putative regulator YpeB. Here, we provide biochemical and genetic evidence that YpeB interacts with and inhibits SleB during dormancy. These findings lay the groundwork for elucidating how SleB is activated during germination.

## INTRODUCTION

Dormant spores can resist extreme environmental insults, including heat, irradiation, desiccation, and antibiotics (1). Although the mechanistic bases for these resistance properties are incompletely understood, it has been appreciated for >50 years that dehydration of the spore core is critical for wet heat resistance (2, 3). During the late stages of sporulation, the spore-specific metabolite pyridine-2,6-dicarboxylic acid (dipicolinic acid or DPA) is synthesized in the mother cell that nurtures the developing spore and is then transported across the double membrane that separates the two cells (4–7). DPA accumulates to very high levels as a Ca^2+^ chelate in the dormant spore and is thought to displace water, contributing to core dehydration and heat resistance (8–10). To maintain this dehydrated state, a thick layer of specialized peptidoglycan (PG) called the cortex is assembled around the spore that physically prevents its expansion by osmosis (11). Sandwiched between the cortex and the spore membrane is a thinner layer of PG, called the germ cell wall, that resembles vegetative PG and serves as the foundational layer for cell wall synthesis during spore outgrowth.

The cortex PG is essential for core dehydration and the maintenance of spore resistance during dormancy (11, 12). Its degradation is also essential for germination and outgrowth (13, 14). In *Bacillus subtilis*, two functionally redundant cortex lytic enzymes, SleB and CwlJ, are responsible for its dissolution during germination (15, 16). Because these enzymes cleave PG at cortex-specific muramic-δ-lactam modifications, they specifically degrade the cortex while sparing the underlying germ cell wall (11).

Bacterial endospores are metabolically inactive and can remain dormant for decades. Yet, In response to nutrient signals, they exit dormancy within minutes and resume growth (13, 14). Accordingly, all the factors required for spore germination are deposited in the spore during its development, including the cortex lytic enzymes (15, 17). Since dormancy can last for decades, it is critical that these lytic enzymes are prevented from cleaving their substrate. How these enzymes are held inactive in the dormant spore is incompletely understood. CwlJ is produced in the mother cell during sporulation and is assembled into the protein coat that surrounds the outer spore membrane (15, 18). CwlJ is therefore physically separated from the cortex layer that is synthesized between the inner and outer spore membranes (11). The outer spore membrane is thought to lose barrier function in the dormant spore (19–21), enabling CwlJ to access the cortex during germination. Importantly, addition of exogenous DPA to dormant spores triggers CwlJ-dependent cortex degradation, suggesting that CwlJ is activated by this spore specific molecule (22, 23). Although this has yet to be demonstrated biochemically, if correct, it provides a mechanism to ensure that CwlJ remains inactive during dormancy. Moreover, because DPA is released from the spore core at the onset of germination (24), it provides a mechanism for CwlJ activation during the exit from dormancy. By contrast, less is known about how SleB is regulated. SleB is produced in the developing spore and is secreted across the inner spore membrane and therefore resides in the same compartment as the cortex (17, 25). In many endospore forming bacteria, SleB is encoded in an operon with the *ypeB* gene (16, 17, 26). YpeB encodes a type I membrane protein with 3 extracytoplasmic PepSY (<u>pep</u>tidase and *B. <u>s</u>ubtilis* <u>Y</u>peB) domains (PF03413) (27, 28). PepSY domains are found on M4 peptidases and have been implicated in regulating their activity (28–30). In the context of this model, YpeB could hold SleB inactive at the inner spore membrane during dormancy and release it during germination.

A recent study reported that AlphaFold3 (AF3) predicts that YpeB and SleB interact and provided genetic evidence consistent with this model (30). Here, we extend these findings, by demonstrating that the two proteins can be co-purified when expressed in *E. coli*. Furthermore, we systematically test the AF3 SleB-YpeB model by mutating residues at the predicted interprotein interface and quantifying protein levels in *B. subtilis* spores. We identify a series of amino acid substitutions in SleB that destabilize YpeB and, reciprocally, substitutions in YpeB that destabilize SleB. Using a subset of these variants, we selected for germination-competent suppressors and identified both general and allele-specific suppressors in the partner protein. In all cases, the suppressor mutants restored the stability of both SleB and YpeB. Collectively, these data provide strong support for the model that YpeB holds SleB inactive in the dormant spore preventing inappropriate cortex degradation during dormancy.

## RESULTS

### SleB and YpeB form a membrane complex

To explore the model that YpeB holds SleB inactive in dormant spores, we used computational structure predictions. Both AlphaFold2 (AF2) and AF3 predicted high-confidence interactions between SleB and YpeB (31, 32). A SleB_2_-YpeB_2_ tetramer yielded even higher confidence metrics (**Fig. 1B and S1**). In these models, each YpeB protomer can be divided into three distinct structural segments (**Fig. 1**). At the N-terminus is a single transmembrane (TM) helix. The TM helix extends unbroken into the extractytoplasmic space, where it participates in a bundle of five helices that we term the helical bundle domain (HBD). A long disordered linker connects the HBD to a C-terminal segment containing three PepSY domains, the first two of which (PepSY1 and PepSY2) are arranged in a C2-pseudosymmetric dimer that forms a single extended β-sheet, and the third of which (PepSY3) forms only a small asymmetric β interaction with PepSY2. PepSY3 bends back to form contacts with the HBD. The two YpeB protomers interact with each other both through their TM helices and their HBDs (**Fig. 1A**). Each SleB catalytic domain is clasped between the PepSY domains and the HBD of one YpeB protomer (**Fig. 1A and 1C**). The PG binding domain of SleB, which is connected to the catalytic domain by a disordered linker, interacts with the outward-facing side of PepSY2 and PepSY3. Interestingly, SleB’s catalytic groove (E203, R222, Y278, H301) is unobstructed but points toward the spore inner membrane (**Fig. 1C**). We propose that, during dormancy, cortex degradation is avoided not by occluding SleB’s active site but rather by holding SleB in a position where it cannot access its substrate.

**Figure 1.**
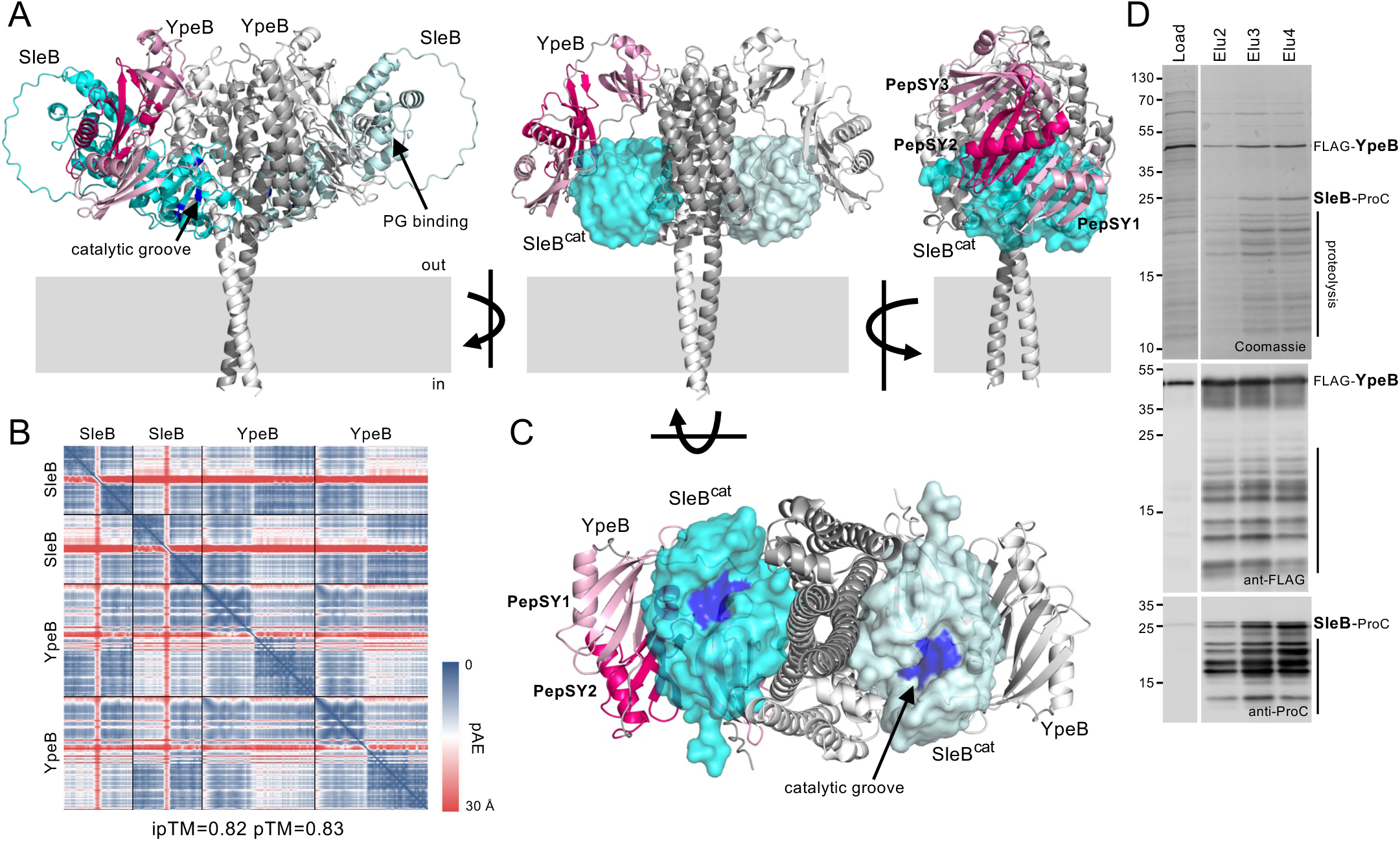
SleB and YpeB assemble into a membrane complex. **(A)** AlphaFold3-predicted structure of the SleB_2_-YpeB_2_ tetrameric complex. YpeB’s PepSY domains embrace SleB’s catalytic domain (SleB^cat^). The individual PepSY domains of one of the YpeB promoters are colored in different shades of pink for clarity. SleB’s peptidoglycan (PG) binding domain and catalytic groove are indicated. The middle and right models only depict SleB’s catalytic domain. **(B)** Plot of the predicted alignment error (pAE) in Å of all residues against all residues for the top-ranked model. Low error (blue) corresponds to well-defined relative domain positions. The template modeling score (pTM) and interface predicted template modeling score (ipTM) for the model are shown below. **(C)** The predicted structure of SleB_2_-YpeB_2_ complex as viewed from inside the spore. The catalytic grooves of SleB’s catalytic domains are shown in blue. **(D)** Representative Coomassie-stained gel and immunoblots of co-purification of SleB-ProteinC (SleB-ProC) and FLAG-YpeB from a detergent-solubilized membrane preparation of *E. coli* cells engineered to express SleB-ProC and FLAG-YpeB. Load and elutions (Elu) from anti-ProC resin are shown. Molecular weight markers in kDa are shown on the left.

Previous immunoblotting experiments determined that SleB and YpeB depend on each other for stability in spores (33). Experimental evidence in support of an interaction between the SleB and YpeB, beyond their dependence on each other for stability in spores, comes from experiments in which we expressed a YFP-YpeB fluorescent fusion in exponentially growing cells with and without co-expression of SleB (**Fig. S2**). When expressed in the absence of SleB, YFP-YpeB accumulated to low levels and displayed weak fluorescence with an occasional fluorescent focus. By contrast, co-expression of SleB resulted in higher levels of YFP-YpeB and multiple fluorescent foci in virtually all cells examined. Although the nature of the YFP-YpeB fluorescent foci is unclear, these data indicate that SleB influences YpeB levels and localization in vegetatively growing cells, suggesting the two proteins interact.

To more rigorously investigate the model that the two proteins form a membrane complex, we sought to co-express SleB and YpeB in *E. coli* and co-purify them. Initial attempts at co-expressing the two proteins revealed that SleB was virtually undetectable in *E. coli* lysates (**Fig. S3**). To improve its expression, we replaced SleB’s native signal peptide with signal peptides from *Erwinia carotovora* PelB and *E. coli* DsbA. The former is secreted post-translationally and the latter co-translationally (34). As can be seen in Figure S3, DsbA’s signal peptide significantly improved SleB expression. We co-expressed this SleB variant fused to a protein C epitope (SleB-ProC) with FLAG-tagged YpeB. Membrane preparations were solubilized with n-dodecyl-D-maltoside and purified FLAG-YpeB on anti-FLAG resin (see Methods). As can be seen in the Coomassie-stained gel and the immunoblots in Figure 1D, SleB-ProC co-purified with FLAG-YpeB. We conclude that the two proteins form a membrane complex, consistent with the AF predictions.

### SleB mutations that decrease SleB/YpeB levels

To probe the AF-predicted complex, we identified a set of amino acid residues in SleB that contact YpeB in the model. We generated amino acid substitutions in these residues in the context of the *sleB-ypeB* locus and introduced the mutated operon at a neutral chromosomal locus in a strain lacking the native *sleB-ypeB* operon. To sensitize the strain and enhance phenotypes of SleB dysfunction, we also deleted *cwlJ*, encoding the functionally redundant cortex lytic enzyme. We sporulated the cells for 30 h and then incubated the sporulation culture at 80 °C for 20 min to kill vegetative cells. The cultures were serially diluted and plated on LB agar plates and the colony forming units were counted. Because SleB stability depends on its interaction with YpeB, amino acid substitutions that disrupt the predict complex will lack cortex lytic enzymes. These spores will fail to germinate due to their inability to degrade the cortex and initiate outgrowth. To maintain standard terminology, we refer to the reduction in colony forming units as a reduction in sporulation efficiency, even though the defect is in germination and outgrowth. Most of the mutants tested displayed a modest (5– to 10-fold) reduction in sporulation efficiency (**Fig. 2B**). However, three mutants, SleB(F237A), SleB(A181L), and SleB(A233L), each caused a >100-fold reduction in sporulation efficiency. To investigate the levels of SleB and YpeB, we analyzed lysates from the purified spores by immunoblot. As anticipated, the mutant spores with reduced sporulation efficiency had very low or undetectable levels of the SleB variants and YpeB, consistent with a failure of the proteins to interact (**Fig. 2C**).

**Figure 2.**
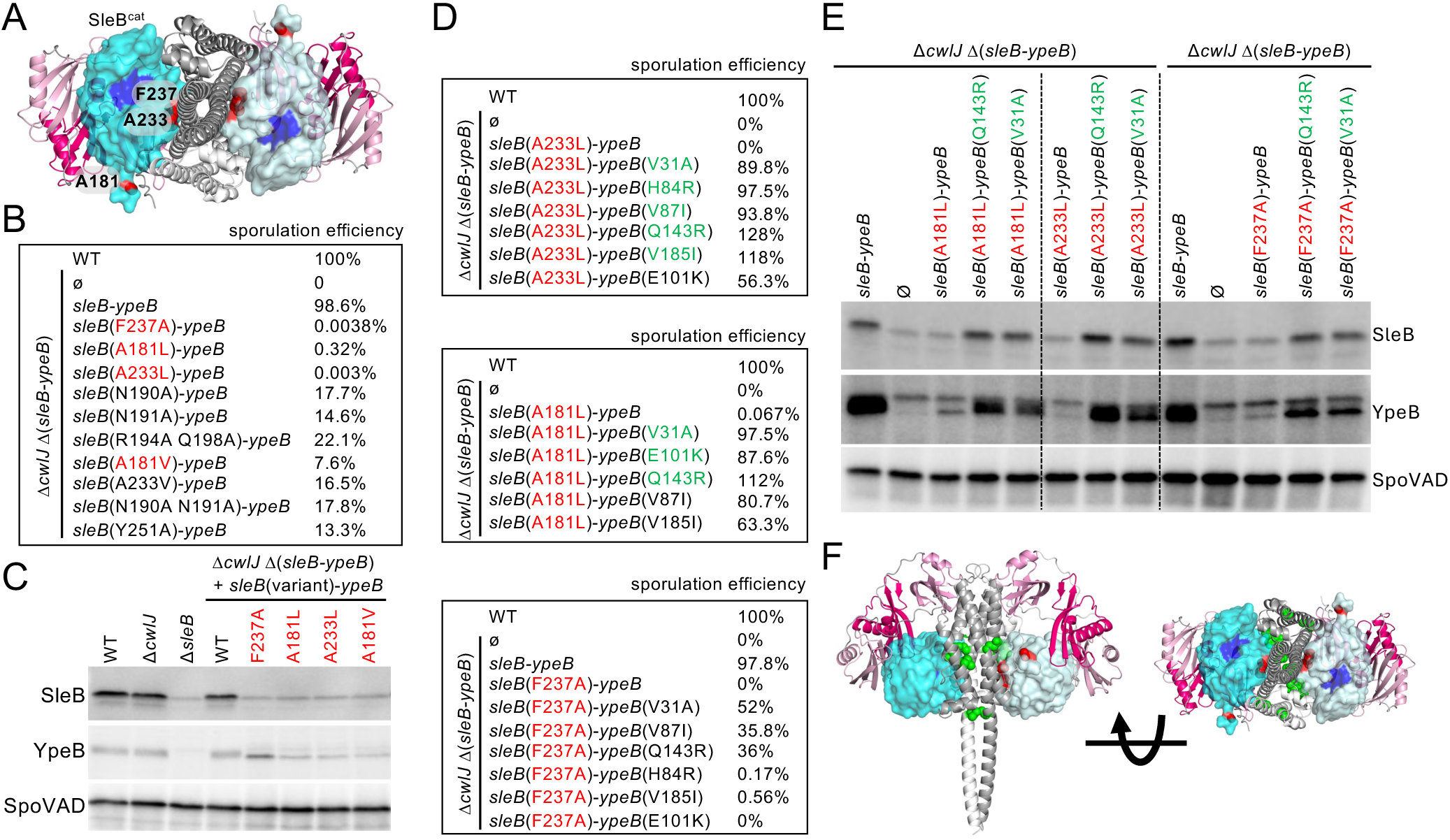
Amino acid substitutions in SleB that disrupt the interaction with YpeB and suppressors that restore it. **(A)** AlphaFold3-predicted structure of the SleB_2_-YpeB_2_ tetrameric complex as viewed from inside the spore. Only the catalytic domains of SleB (SleB^cat^) are shown. The catalytic grooves (blue) and the three amino acid residues (red) that are important for interaction with YpeB are highlighted. **(B)** Sporulation efficiencies of the indicated SleB mutants. **(C)** Representative immunoblots of SleB and YpeB from spore lysates of the indicated strains. SpoVAD controls for loading. **(D)** Sporulation efficiencies of the indicated SleB mutants and YpeB suppressors. Suppressors mutations shown in green were identified in the enrichment screen with the indicated SleB mutant. Suppressors shown in black were identified in one of the other enrichment screens but were tested for general suppression. **(E)** Representative immunoblots of SleB and YpeB from spore lysates of the indicated strains. SpoVAD controls for loading. **(F)** AlphaFold3-predicted structure of the SleB_2_-YpeB_2_ tetrameric complex from side and below. Residues in SleB (A181 A233, and F237) that disrupt interaction are shown in red. YpeB suppressors that suppress the sporulation defect and restore protein stability are shown in green.

The identification of predicted interaction interface *sleB* mutations that cause YpeB and SleB instability provides support for the AF model. However, we could not exclude the possibility that the loss of SleB was unrelated to its interaction with YpeB. If the instability of the SleB variants was indeed due to a failure to interact with YpeB, we reasoned we should be able to identify mutations in *ypeB* that suppress the sporulation defect. Accordingly, we PCR-mutagenized *ypeB* in the presence of the *sleB* mutation and selected for suppressors that could form germination-competent, heat-resistant spores (see Methods). We identified five *ypeB* suppressors of *sleB*(A233L) and three *ypeB* suppressors of *sleB*(A181L) (**Fig. 2D**). Interestingly, two of the suppressor mutations (V31A and Q143R) were isolated in both selections, suggesting that these are general rather than allele-specific suppressors. Consistent with this idea, neither V31 nor Q143 in YpeB are in close proximity to A233 or A181 in SleB in the AF model (**Fig. 2F**). In fact, none of the amino acid substitutions in the YpeB suppressors was predicted to be in close proximity to the original substitutions in SleB (**Fig. 2F**). We tested several of the *ypeB* suppressors identified in the *sleB*(A233L) selection for their ability to suppress *sleB*(A181L) and separately *sleB*(F237A). Most restored sporulation efficiency (**Fig. 2D**) and germination properties (**Fig. S4**), as assayed by change in optical density, to near wild-type levels, indicating that they are general suppressors. Finally, we investigated the levels of SleB and YpeB in spores harboring the *sleB* interaction mutation with and without the *ypeB* suppressors. As can be seen in the immunoblots in Figure 2E, in all cases analyzed, the suppressors restored the levels of both SleB and YpeB. These results argue that the original amino acid substitutions in SleB (A181L, A233L, and F237A) impaired or abolished the ability of SleB to interact with YpeB and therefore provide support for the AF-predicted SleB-YpeB model. Furthermore, the isolation of *ypeB* suppressors of these *sleB* mutations provides genetic evidence for an interaction between the two proteins.

### YpeB mutations that decrease SleB/YpeB levels

Next, we sought to systematically test amino acid residues in YpeB that were predicted to interact with SleB. We targeted 16 YpeB residues (**Fig. 3A**), generating single, double, and triple mutations. Consistent with the large AF-predicted interface, 13 of the 16 single mutations had modest or undetectable sporulation defects (**Fig. 3B**). Furthermore, three double mutations and one triple mutation had similarly modest sporulation defects. However, three *ypeB* single mutations (E444R, R147E, and N37A) and two double mutations had >1000-fold reduction in sporulation efficiency (**Fig. 3B**). In virtually all cases, the levels of SleB and YpeB in the mutants correlated with sporulation efficiency (**Fig. 3C**) and germination properties (**Fig. S5**).

**Figure 3.**
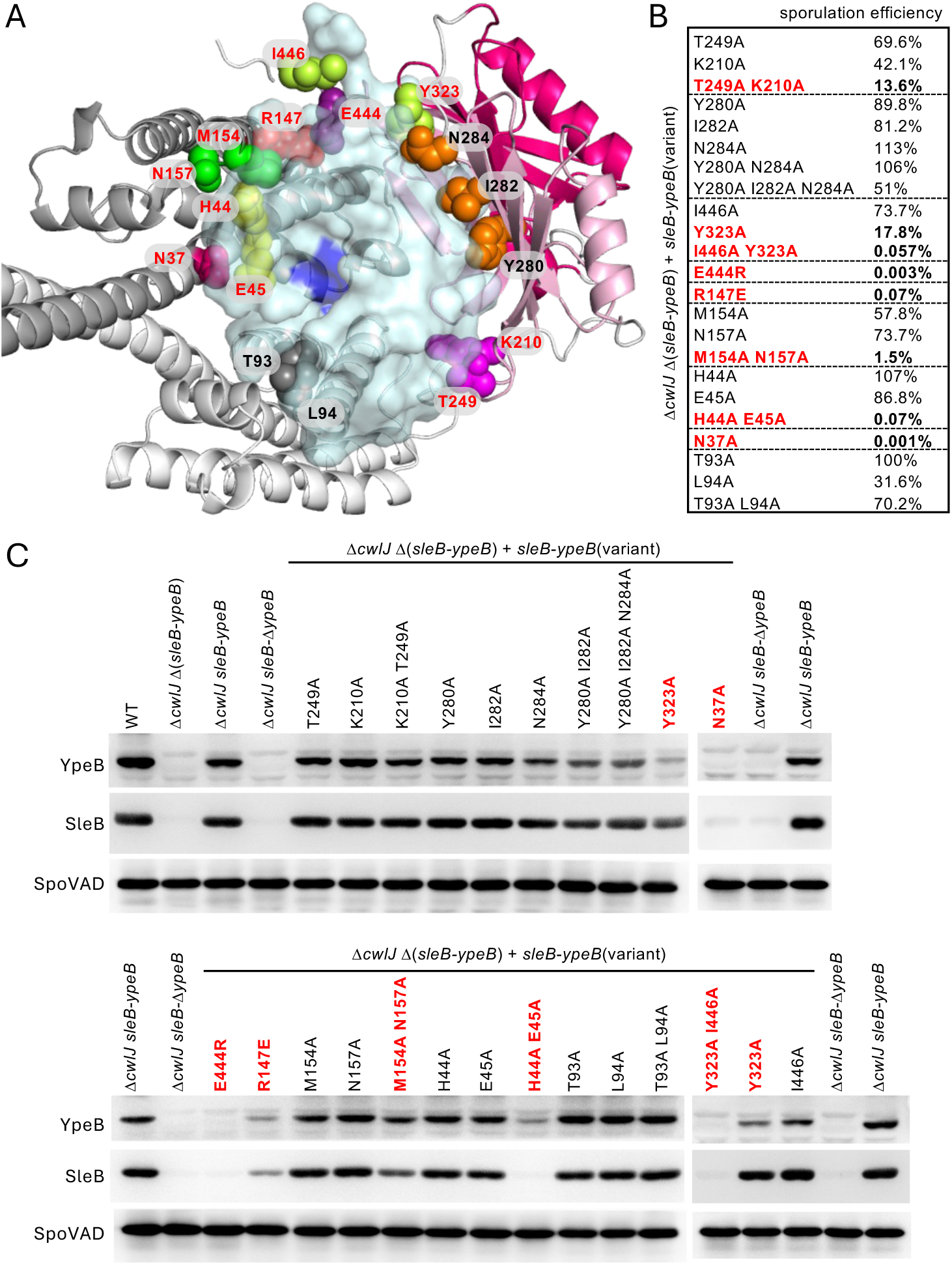
Amino acid substitutions in YpeB that disrupt the interaction with SleB. **(A)** AlphaFold-predicted structure of the YpeB dimer (dark/light grey) bound to the catalytic domain of SleB (cyan) viewed from the spore membrane. YpeB residues that are predicted to interact with SleB are shown as colored spheres. **(B)** Sporulation efficiencies of the indicated YpeB variants in a Δ*cwlJ* background. **(C)** Representative immunoblots of spore lysates from the indicated strains probed for YpeB and SleB. SpoVAD controls for loading. The blots on the right were from a different set of gels, but the spore lysates were generated and analyzed at the same time as those on the left. YpeB variants highlighted in red are impaired in sporulation efficiency.

### Identification of general and allele-specific suppressors

While the identification of destabilizing *sleB* mutations and general *ypeB* suppressors provides support for a direct interaction between the two proteins, we sought to identify allele-specific *sleB* suppressors of two strong *ypeB* interaction mutations, *ypeB*(R147E) and *ypeB*(E444R). We PCR-mutagenized *sleB* in the presence of the *ypeB* mutations and selected for suppressors that could form heat resistant spores that were germination-competent. We identified *sleB*(D192V) as a strong suppressor of *ypeB*(R147E) (**Fig. 4C**). The suppressor dramatically improved sporulation efficiency, restored wild-type germination kinetics (**Fig. 4E**), and stabilized both YpeB(R147E) and SleB(D192V) (**Fig. 4D**). We then combined the *sleB*(D192V) mutation with several other *ypeB* interaction mutations and tested the double mutants for sporulation efficiency. Only *ypeB*(R147E) was suppressed by *sleB*(D192V), establishing this pairing as an allele-specific interaction and providing support for the AF-predicted interface structure in which these two residues are in physical proximity (**Fig. 4A**). The chemical environment of this region of the predicted interface is also consistent with the suppression result: YpeB-R147 lies near a cluster of three acidic residues (YpeB-E444, YpeB-D51, SleB-D192). Increasing the negative charge by –2 with the *ypeB*(R147E) mutation likely creates unfavorable repulsions, while removing one of those negative charges with the *sleB*(D192V) could relieve it.

**Figure 4.**
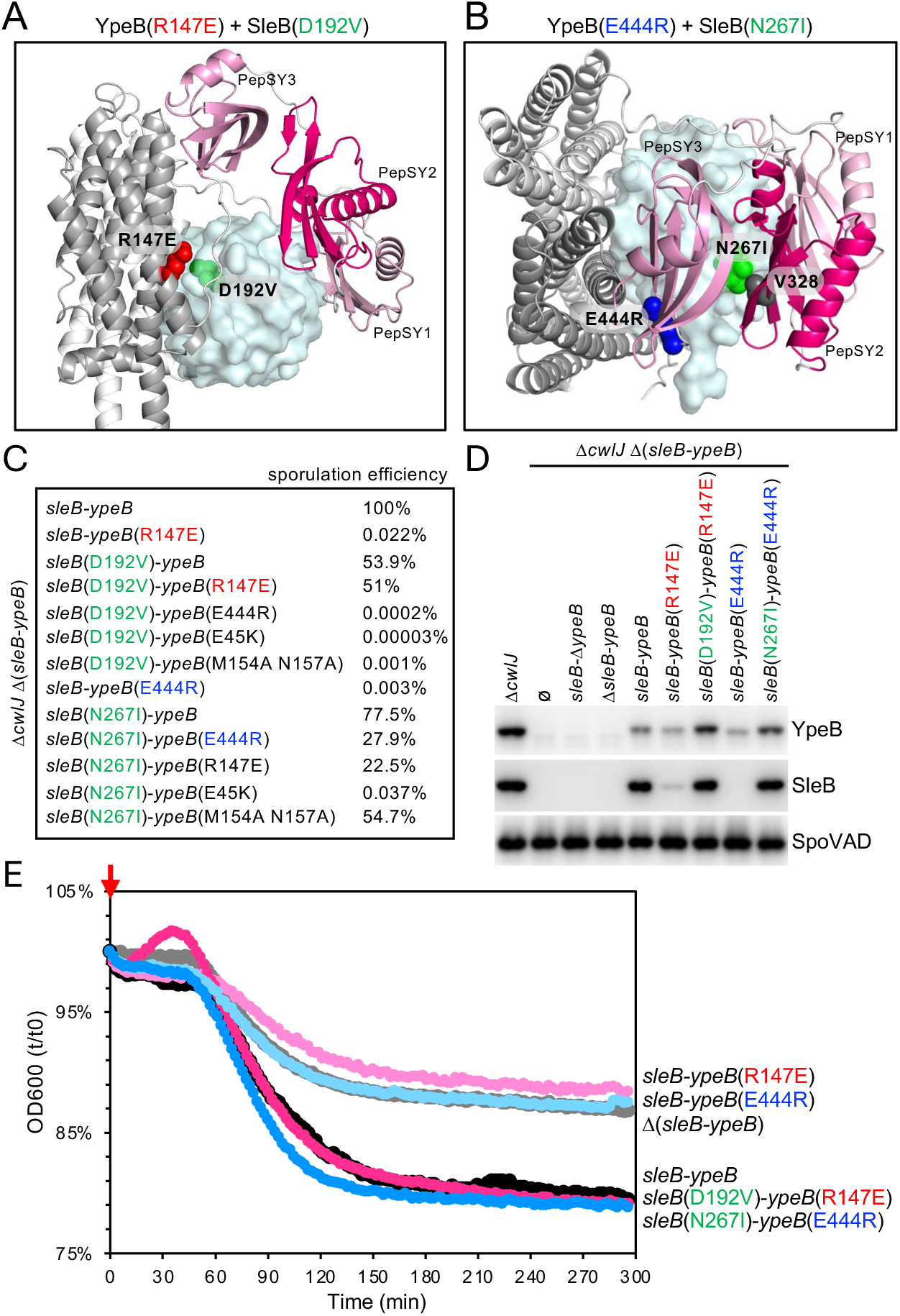
Identification of an allele-specific suppressor of *ypeB*(R147E). **(A)** AlphaFold3-predicted structures of the YpeB dimer (light/dark grey) and the catalytic domain of SleB (cyan). YpeB’s PePSY domains are in shades of pink. The R147E substitution in YpeB (red) and the D192V substitution in SleB (green) that suppresses the YpeB mutant are indicated. **(B)** The YpeB(E444R) mutant (blue) and its suppressor SleB(N267I) (green) are indicated. Valine-328 (V328) in SleB’s PepSY2 domain that is hypothesized to interact with N267I is shown in dark grey. **(C)** Table of sporulation efficiencies of the indicated SleB and YpeB mutants. **(D)** Representative immunoblots from spore lysates of the indicated strains probed for SleB and YpeB. SpoVAD controls for loading. **(E)** Representative graph of spore germination in response to L-alanine, as assayed by a reduction in optical density (OD_600_). Histodenz-purified phase-bright spores of the indicated strains lacking *cwlJ* were induced to germinate with 10 mM L-alanine (red arrow), and the OD_600_ was monitored over time. The data are plotted as the percent reduction in OD_600_ relative to time 0.

We also identified a *sleB* suppressor (N267I) of *ypeB*(E444R) (**Fig. 4C**). This suppressor restored sporulation efficiency, germination kinetics, and YpeB(E444R) stability (**Fig. 4D-E**). However, N267 was not predicted to reside in close proximity to YpeB-E444 (**Fig. 4B**) and the *sleB* mutation suppressed several strong *ypeB* interaction mutations, consistent with the idea that N267I improved the interaction between SleB and YpeB in a manner that is independent of the E444R mutation (**Fig. 4C**). Interestingly, N267 resides in close proximity to V328 in YpeB’s PepSY2 domain (**Fig. 4B**). Accordingly, one possible explanation for the general suppression by N267I is the strengthening of the interaction between PepSY2 and SleB through increased hydrophobic contacts.

## DISCUSSION

Altogether, our findings indicate that *B. subtilis* YpeB and SleB reside in a membrane complex and provide strong support for the AF models that YpeB holds SleB inactive. In this model, a YpeB dimer binds two SleB molecules. The three PepSY domains from each YpeB protomer embrace a SleB catalytic domain and position its catalytic groove toward the spore membrane and away from the cortex peptidoglycan. Importantly, AF3 predicts similar high-confidence interactions between YpeB and SleB from a diverse set of endospore-forming Bacillota (**Fig. S6**). These findings along with the work on *Bacillus cereus* SleB-YpeB from Christie and co-workers (30) establish that YpeB is required to hold SleB inactive in dormant endospores and provides a framework for investigating how YpeB releases SleB during germination.

Although we could provide a plausible mechanism to explain how SleB(N267I) functions as a general suppressor, it is less clear how the YpeB suppressors identified function to suppress the three SleB interaction interface mutants A181L, A233L, F237A. Two of the YpeB suppressors (H84R and E101K) are predicted to contact SleB and could stabilize the YpeB-SleB interaction, as we have proposed for SleB(N267I). However, four YpeB suppressors (V31A, V87I, Q143R, V185I) are not predicted to make direct contact with SleB. In these cases, we hypothesize that these amino acid substitutions stabilize the global YpeB fold and this could be sufficient to stabilize its interaction with SleB. Mechanistic understanding of these suppressors awaits experimental structure determination of the YpeB-SleB complex.

### SleB instability as a central feature of dormancy

Previous studies demonstrating that SleB and YpeB depend on each other for stability (33) motivated the hypothesis that YpeB interacts with SleB and holds it inactive. Our biochemical data establishing that the two proteins reside in a complex and our genetic validation of the AF models prompt us to interpret SleB’s instability in a broader biological context. We propose that degradation of SleB molecules that fail to interact with YpeB upon secretion into the intermembrane space is an evolved feature of spore maturation and dormancy. SleB’s instability ensures that this potent lytic enzyme is not packaged in the dormant spore adjacent to its substrate free of its inhibitor.

### A potential role for YpeB’s PepSY domains as a chaperone-inhibitor

To further explore the role of YpeB’s regulation of SleB, we examined the role of the PepSY domain on the secreted M4 peptidase family (28). Extracellular microbial proteases, including those in the M4 family, are often synthesized as pro-enzymes (35). In many cases, the pro-peptide has been shown to function as an intramolecular chaperone-inhibitor (36, 37). In Gram-negative bacteria, these domains are essential for folding the protease upon secretion across the cytoplasmic membrane (38–40). After enzyme maturation, the protease undergoes autoproteolysis, cleaving its pro-domain (35, 37). However, the pro-peptide remains non-covalently associated with the mature protease and functions to inhibit proteolytic activity until it is secreted across the outer membrane and released into the environment. The pro-domains of M4 peptidases contain a PepSY domain and a fungalysin/thermolysin propeptide (FTP) domain (PF07504) that together function as chaperone and inhibitor (39). Mutational analysis of the pro-domain of the M4 peptidase LasB in *Pseudomonas aeruginosa* suggests that the PepSY domain contributes to autoinhibition of LasB’s metallopeptidase activity (29). Whether the PepSY domain also contributes to chaperone activity has not been rigorously investigated. Intriguingly, the PepSY and FTP domains on M4 peptidases like LasB and M36 peptidases like fungalysin both form beta-sheets that are buttressed by alpha-helices and together the two domains embrace the peptidase domain in a manner that is reminiscent of the embrace of SleB by YpeB’s PepSY domains (**Fig. S7**).

Given SleB’s instability in the absence of YpeB, we propose that YpeB’s PepSY domains possess both chaperone and inhibitory activities. In the context of this model, SleB molecules that fail to interact with YpeB remain partially or fully unfolded in the intramembrane space and are subject to proteolytic degradation during spore maturation. Degradation ensures that these lytic enzymes do not have the opportunity to fold during prolonged periods of dormancy and therefore cannot inappropriately degrade the cortex. Interestingly, we have found that expression of SleB and YpeB from distinct chromosomal loci results in reduced levels of both proteins in dormant spores (**Fig. S8**), suggesting that YpeB’s proposed chaperone activity is required during secretion of SleB into the intermembrane space. Once folded, SleB remains stably bound and inhibited by YpeB in the dormant spore. Upon exposure to germinants, the mature enzyme is released by a mechanism that remains to be elucidated.

## MATERIALS AND METHODS

### General methods

All strains used in this study were derived from *Bacillus subtilis* 168 (41). All deletion mutants were derived from the *Bacillus* knock-out collection (BKE) (42), or constructed by direct transformation of isothermal assembly products. Antibiotic cassettes were excised using a temperature-sensitive plasmid that constitutively expresses Cre recombinase (Meeske). Site-directed mutants were generated using a modified QuickChange protocol. Sporulation was induced by nutrient exhaustion in Difco Sporulation Medium (DSM) at 37 °C for 30h (43). All plasmids, strains and primers used in this study, can be found in supplementary information. Experiments presented in the text and supplementary figures, were from one of at least three biological replicates.

### Spore purification

Spores used for germination assays or immunoblot analysis were generated by sporulating a lawn of cells on DSM agar plates at 30 °C for 96 h. Spores were scraped from the plates, washed 3 times with ddH_2_O, and then resuspended in 350 µL 20% histodenz (Sigma-Aldrich). The suspension was layered on top of 1 mL 50% histodenz in a microfuge tube and the step-gradient was centrifuged at 15K rpm for 30 min at 4 °C. The pellet fraction containing the phase-bright spores was collected and washed 6X with ddH_2_O. Spores were stored at 4 °C and analyzed within 72 h. Purified phase-bright spores were >95% pure based on phase-contrast microscopy.

### Germination assay using reduction in optical density

Purified phase-bright spores were normalized to an OD_600_ of 1.2 in 25 mM HEPES pH 7.4, and heat-activated at 70 °C for 30 min, followed by incubation on ice for 15 min. 100 µL spore resuspension were transferred to a clear, flat-bottom, 96-well plate. An equal volume of 25 mM HEPES pH 7.4 (Buffer) or 10 mM L-Ala resuspended in 25 mM HEPES pH 7.4, was added to the spore suspensions to a final OD_600_ of 0.6. The OD_600_ was monitored every 2 min for 5 h using an Infinite M plex plate reader (Tecan). The plate was maintained at 30 °C with agitation between measurements. All samples were analyzed in technical triplicate.

### Co-expression and co-purification of *B. subtilis* SleB and YpeB in *E. coli*

Plasmid pYG566, a pET-DUET derivative for co-expression of SP^DsbA^-SleB-ProC and His-SUMO-FLAG-YpeB was transformed into *E. coli BL21*(DE3) carrying pAM174 (44). pAM174 is an arabinose-inducible expression plasmid used to express Ulp1 under arabinose control. Transformants were grown in Terrific Broth supplemented with 0.4% glycerol. 0.1% glucose, 2 mM MgCl_2_, 100 μg/mL ampicillin and 20 μg/mL chloramphenicol, at 37 °C with agitation. When the culture reached an OD_600_ of ∼0.2, it was transferred to 20 °C shaker-incubator. When the OD_600_ reached 0.7, IPTG (0.5 mM IPTG, final) and arabinose (0.1%, final) were added. After an 18 h induction, cells were harvested by centrifugation at 8000 rpm for 15 min at 4 °C. The cell pellet was resuspended in 45 mL lysis buffer (50 mM HEPES at pH 7.6, 150 mM NaCl, 20 mM MgCl_2_ and 0.5 mM DTT) supplemented with 5 U benzonase (Sigma) and 1X complete protease inhibitor (Roche), and lysed by two passages in a cell disrupter (Constant System) at 25,000 psi. The lysate was subjected to ultracentrifugation at 35K rpm for 1 h at 4 °C, and the membrane pellet fraction was homogenized and solubilized in 50 mL Buffer H (20mM HEPES at pH 7.6, 150 mM NaCl, and 20% glycerol) containing 1% n-dodecyl-β-D-maltopyranoside (DDM, Thermo Fisher), and rotated end-over-end at 4 °C for 2h. The detergent-solubilized membrane fraction was then subjected to ultracentrifugation at 35K rpm at 4 °C for 1h. The supernatant was collected and supplemented with CaCl_2_ (2 mM final). The DDM-solubilized membrane proteins were loaded on an anti-M1 FLAG antibody resin. The resin was washed with 25-column volumes (CVs) of wash buffer (20 mM HEPES at pH 7.6, 150 mM NaCl, 20% glycerol, 2 mM CaCl_2_ and 0.1% DDM), and the bound proteins were eluted with 5 CVs elution buffer [20 mM HEPES at pH 7.6, 150 mM NaCl, 10% glycerol, 5 mM EDTA at pH 8.0, 0.1% DDM and 0.2 mg/mL FLAG peptide (DYKDDDDK, Sigma)]. The eluted protein fractions were resolved by SDS-PAGE on a 17.5% polyacrylamide gel and visualized by InstantBlue Coomassie staining (ISB1L) and by immunoblot as described below.

### Spore lysates and immunoblot analysis

Purified phase-bright spores (OD_600_=30) were resuspended in 500 μL cold PBS supplemented with 1 mM phenylmethylsulfonyl fluoride (PMSF) and transferred to 2 mL tubes containing Lysis Matrix B (MP Biomedicals). Samples were incubated on ice for 15 min and lysed using a FastPrep instrument (MP Biomedicals) at 6.5 m/s for 60 s. Immediately afterwards, an equal volume of 2× sample buffer (250 mM Tris-HCl, pH 6.8, 10 mM EDTA, 4% SDS, and 20% glycerol) supplemented with 10% β-mercaptoethanol was added and the samples were mixed by vortexing. Lysates were clarified by centrifugation at 15,000 rpm for 5 min, and the supernatants were collected. Protein concentrations were determined using a noninterfering protein assay (G-Biosciences), and samples were normalized, resolved by SDS-PAGE on 17.5% polyacrylamide gels, and transferred to PVDF membranes (Immobilon-P, Millipore). The PVDF membranes were blocked with 5% nonfat milk dissolved in 1X PBS with 0.5% Tween-20 (PBST), and then incubated overnight at 4 °C with anti-SleB (1:5,000) (25), anti-YpeB (1:2,500) (25), and anti-SpoVAD (1:10,000) (45) polyclonal antibodies diluted in 3% BSA in PBST. The blots were then washed 4 times with PBST and then incubated with goat anti-rabbit antibody coupled to horseradish peroxidase (Bio-Rad) for 2 h at room temperature. The blots were again washed 4 times and then incubated with Western Lightning ECL reagent (PerkinElmer) and visualized using a Bio-Techne FluorChem R System.

For *E. coli*-derived protein samples, lysates were resolved by SDS-PAGE and transferred as described above. Membranes were probed with anti-ProC (1:1,000) (46), anti-FLAG (1:5,000; Sigma-Aldrich), or anti-SleB antibodies (25) (47), followed by detection with horseradish peroxidase conjugated goat anti-mouse or anti-rabbit secondary antibodies.

### PCR-mutagenesis and suppressor selections

To screen for suppressors of *ypeB(R147E)* and *ypeB(E444R)* the *sleB* gene was PCR-amplified with primers oYG1000 and oYG1001 using an error-prone pfu polymerase (48) using pYG534 harboring the *sleB-ypeB* locus and a *kan* cassette as template. The PCR products were cloned by isothermal assembly into pYG767 [*yhdG::P_sleB_-(HindIII-XhoI)-ypeB(R147E) spec (amp)*] or pYG768 [*yhdG::P_sleB_-(HindIII-XhoI)-ypeB(E444R) spec (amp)*] cut with HindIII and XhoI. The reactions were transformed into *E. coli DH5α*, generating ∼100,000 transformants each. To screen for suppressors of *sleB*(A181L) and *sleB*(A233L), the plasmid libraries were constructed similarly. The *ypeB* gene was PCR-mutagenized with primers oYG1003 and oYG1006, using pYG534 as template, and the mutagenized PCR products were used as megaprimers for amplification using pYG684 and pYG685 as template, followed by restriction with DpnI at 37 °C for 1 h. The DpnI-treated products were micro-dialyzed using 0.025 μm membrane (Millipore) floated on ddH_2_O for 20 min. The dialyzed products were transformed into *E. coli 10-Beta*(NEB) electro-competence cells, resulting in ∼220,000 transformants each.

The four *E. coli* libraries were separately pooled and stored at –80 °C. Plasmids from 20 individual transformants from each library, were isolated and subjected to long-read sequencing (Genewiz). Each library had ∼1.3 mutations per kilobase in the target gene. The plasmid libraries of *sleB*-ypeB(R147E)* and *sleB*-ypeB(E444R)* were transformed into *B. subtilis* strain bYG2282 lacking the native *sleB-ypeB* locus and *cwlJ*, generating ∼60,000 and ∼75,000 transformants, respectively. The plasmid libraries of *sleB(A181L)-ypeB\** and *sleB(A233L)-ypeB\** were transformed into *B. subtilis* strain bYG1978 lacking the native *sleB-ypeB* locus and *cwlJ*, resulting in ∼130,000 and 180,000 transformants, respectively.

To isolate suppressors, each pooled *B. subtilis* library was grown in liquid DSM sporulation medium for 30 h at 37 °C, and the sporulated culture was incubated at 80 °C for 20 min to kill vegetative cells and sporulation-defect mutants. The cultures were then plated on LB agar. Approximately 50,000 heat-resistant colonies were pooled for another round of sporulation, heat-kill, and plating.

After the second round of enrichment, 30 individual heat-resistant colonies from each library, were separately picked and inoculated in DSM medium and sporulated as described above. The sporulation efficiency was determined for each strain and genomic DNA from mutants that had sporulation efficiencies similar to wild-type was prepared. The ectopic mutagenized *sleB-ypeB* locus was PCR-amplified and the product sequenced. All mutants presented in the paper, were re-made by site-direct mutagenesis and re-test for suppression.

### Structural modeling using AlphaFold-Multimer

All predicted structures were generated using Alphafold-multimer-v2 or –v3 and ColabFold (31, 32, 49), run locally on Harvard Medical School O2 computing Cluster or on the AlphaFold Server. General parameters were as follows: multiple sequence alignments (MSA) were built using mmseqs2. Sequences from the same operon were paired in the MSA, and both paired and unpaired sequences were used to generate models. Five models were generated in each run, each model was relaxed using AMBER and ranked by pTM score. The maximum number of model recycling was set to 12 and the top ranked models were used throughout the text.

### Analysis of SleB-YpeB homologs

A local psi-blast run was performed on the amino acid sequence of *B. subtilis* YpeB against the RefSeq Select Protein database using an e-value of 0.05 and 5 iterations. A subset of the YpeB homologs were used to query Gene Neighborhood Analysis (50) to identify operons that contained *sleB* and *ypeB*. 25 diverse SleB/YpeB pairs were chosen for further analysis. A phylogenetic tree was built using the YpeB orthologs to determine the diversity of proteins identified (51). The signal peptide on each SleB homolog was determined using DeepTMHMM (52) and was excluded in follow up analysis. Each SleB-YpeB pair was analyzed for interaction using AlphaFold3 (31) and the pTM and ipTM for the best model was reported in Figure S6. Eight SleB-YpeB pairs that were phylogenetically most distant from each other were selected and predicted structures and pAE plots generated and presented in Figure S6.

## ACKNOWLEDGEMENTS

We thank all members of the Bernhardt-Rudner supergroup for helpful advice, reagents, discussions, and encouragement, Peter Setlow and Anne Moir for antisera, and the MicRoN core for advice on microscopy. Support for this work comes from the National Institute of Health Grants R01AI164647 and R21AI171308 (DZR).

## SUPPPLEMENTAL FIGURE LEGENDS

**Figure S1.**
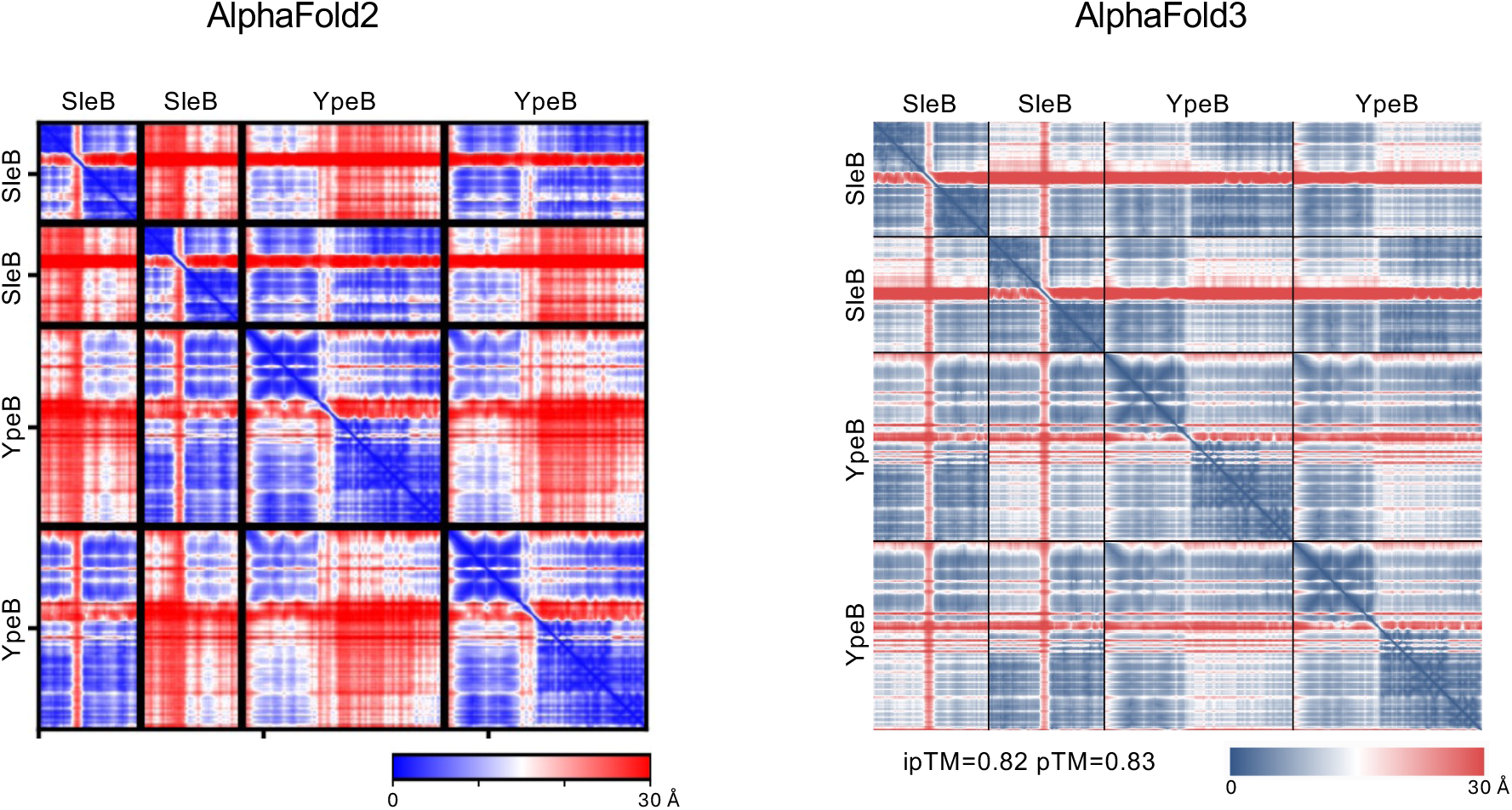
Predicted alignment error (pAE) plots for SleB_2_-YpeB_2_ AlphaFold-Multimer predictions. Plots of the predicted alignment error (pAE) in Å of all residues against all residues for the top-ranked AlphaFold2 and AlphaFold3 models for the SleB_2_-YpeB_2_ tetramer. Low error (blue) corresponds to well-defined relative domain positions. The interface predicted template modeling score (ipTM) and the template modeling score (pTM) for the AlphaFold3 model are shown below the plot.

**Figure S2.**
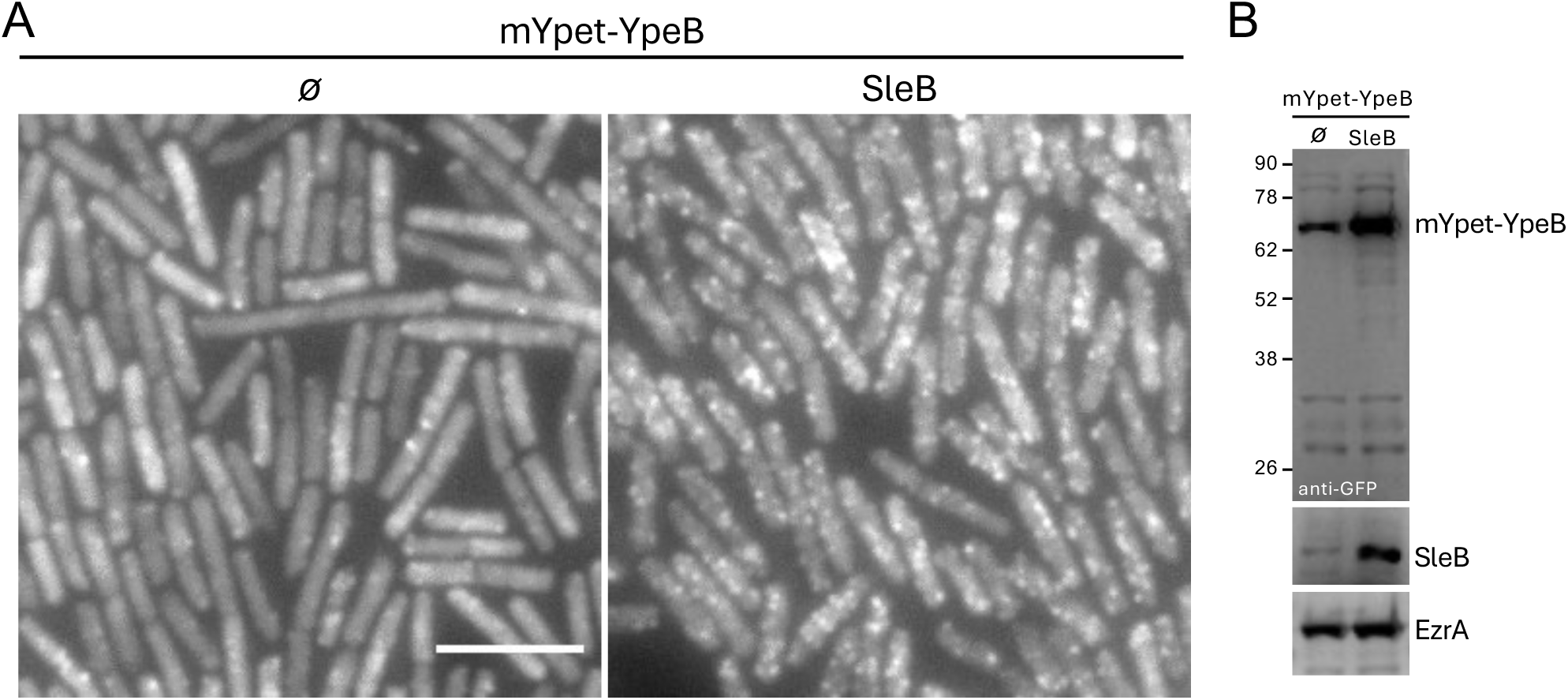
SleB expression stabilizes YpeB and promotes mYpet-YpeB fluorescent foci when expressed during growth. **(A)** Representative fluorescence images of exponentially growing *B. subtilis* cells expressing mYpet-YpeB in the absence (ø) or presence of SleB. The mYpet-YpeB fusion was expressed under the control of a xylose-regulated promoter (PxylA) with 3.3 mM xylose, and SleB was expressed under the control of an IPTG-regulated promoter (Phyperspank) with 1 mM IPTG. Scale bar indicates 5 μm. **(B)** Representative immunoblot of lysates from the strains in (A). mYpet-YpeB and SleB were detected with anti-GFP and anti-SleB antibodies. EzrA controls for loading. Molecular weight standards (in kDa) are shown on the left.

**Figure S3.**
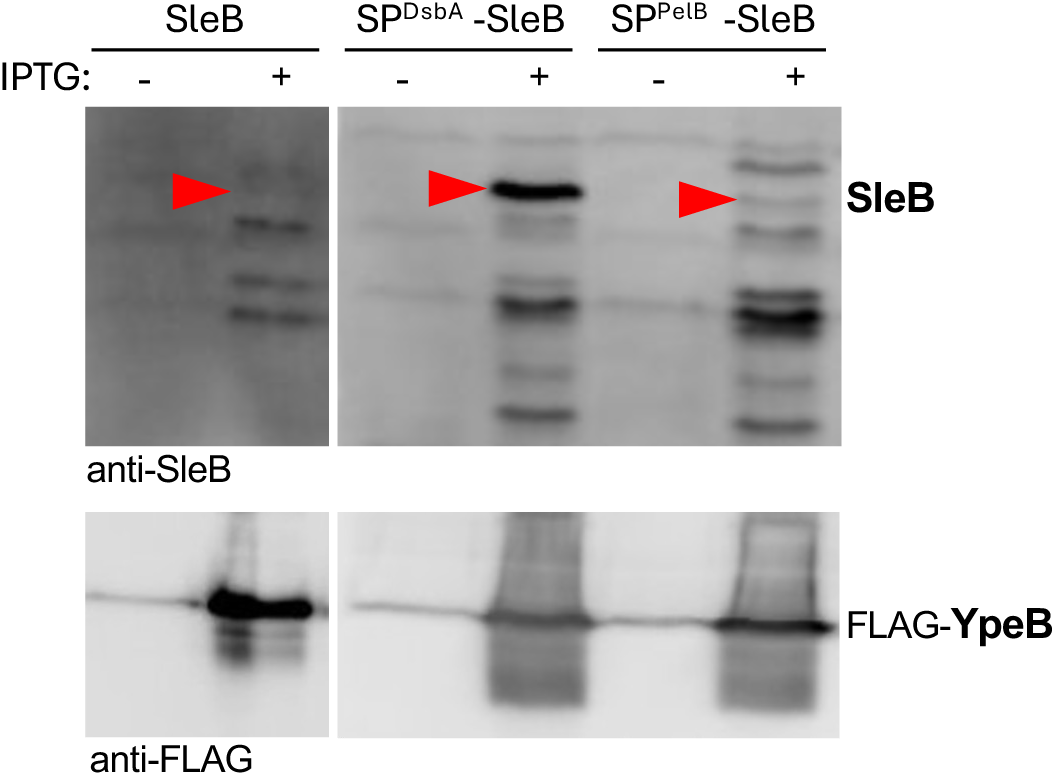
*E. coli* expression of SleB is enhance when its native signal peptide is replaced with *E. coli*’s DsbA signal peptide. Representative immunoblots of SleB and FLAG-YpeB co-expressed in *E. coli* with 0.5 mM IPTG. Cells expressing wild-type *sleB* and *sleB* variants with the signal peptide (SP) from *E. coli* DsbA and from *Erwinia carotovora* PelB are shown. The band corresponding to SleB is indicated (red caret).

**Figure S4.**
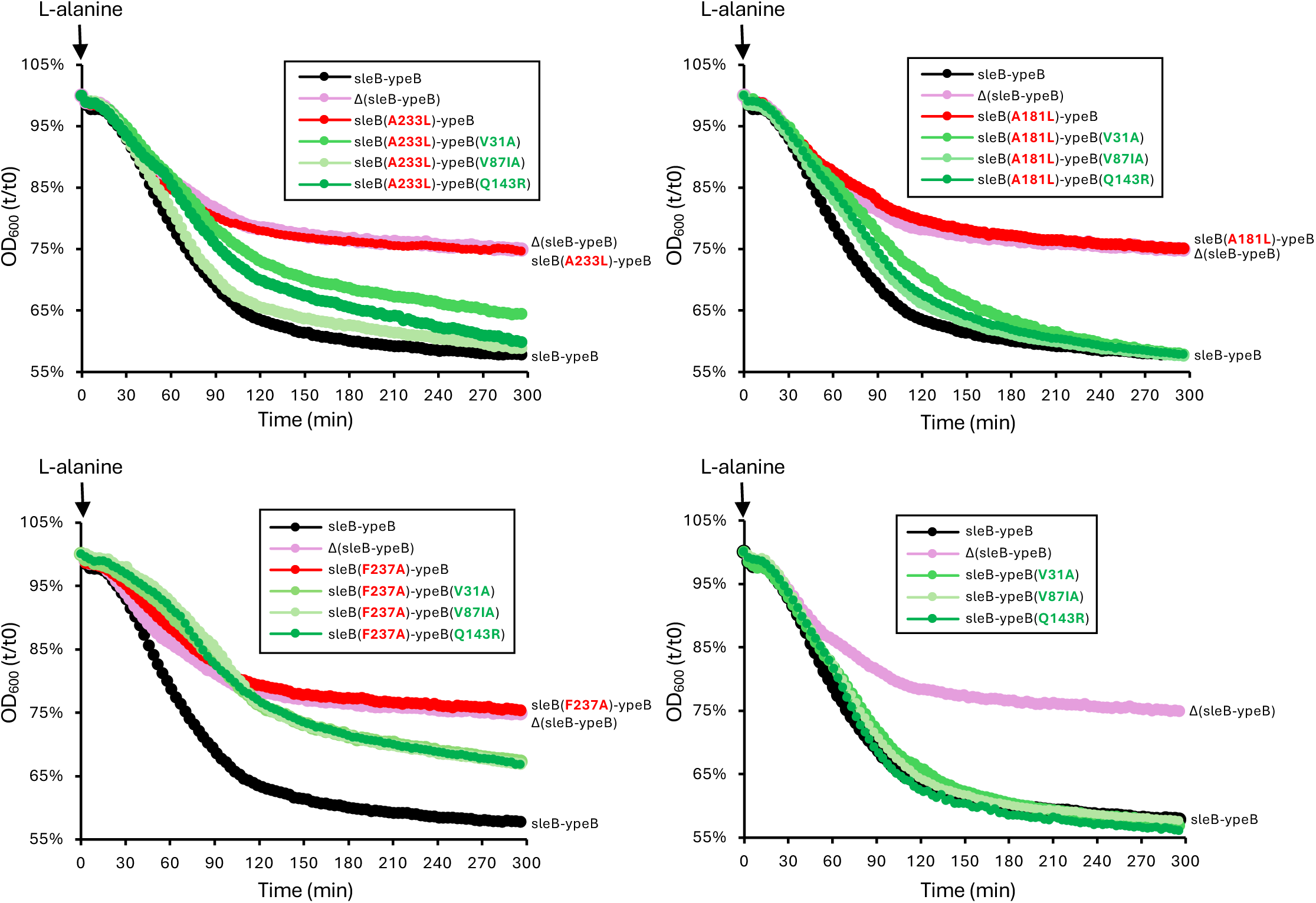
YpeB suppressors mutants are germination-competent. Representative graphs of spore germination in response to L-alanine as assayed by a reduction in optical density (OD_600_). Histodenz-purified phase-bright spores of the indicated strains lacking *cwlJ*, were induced to germinate with 10 mM L-alanine, and OD_600_ was monitored over time. The data are plotted as the percent reduction in OD_600_ relative to time 0.

**Figure S5.**
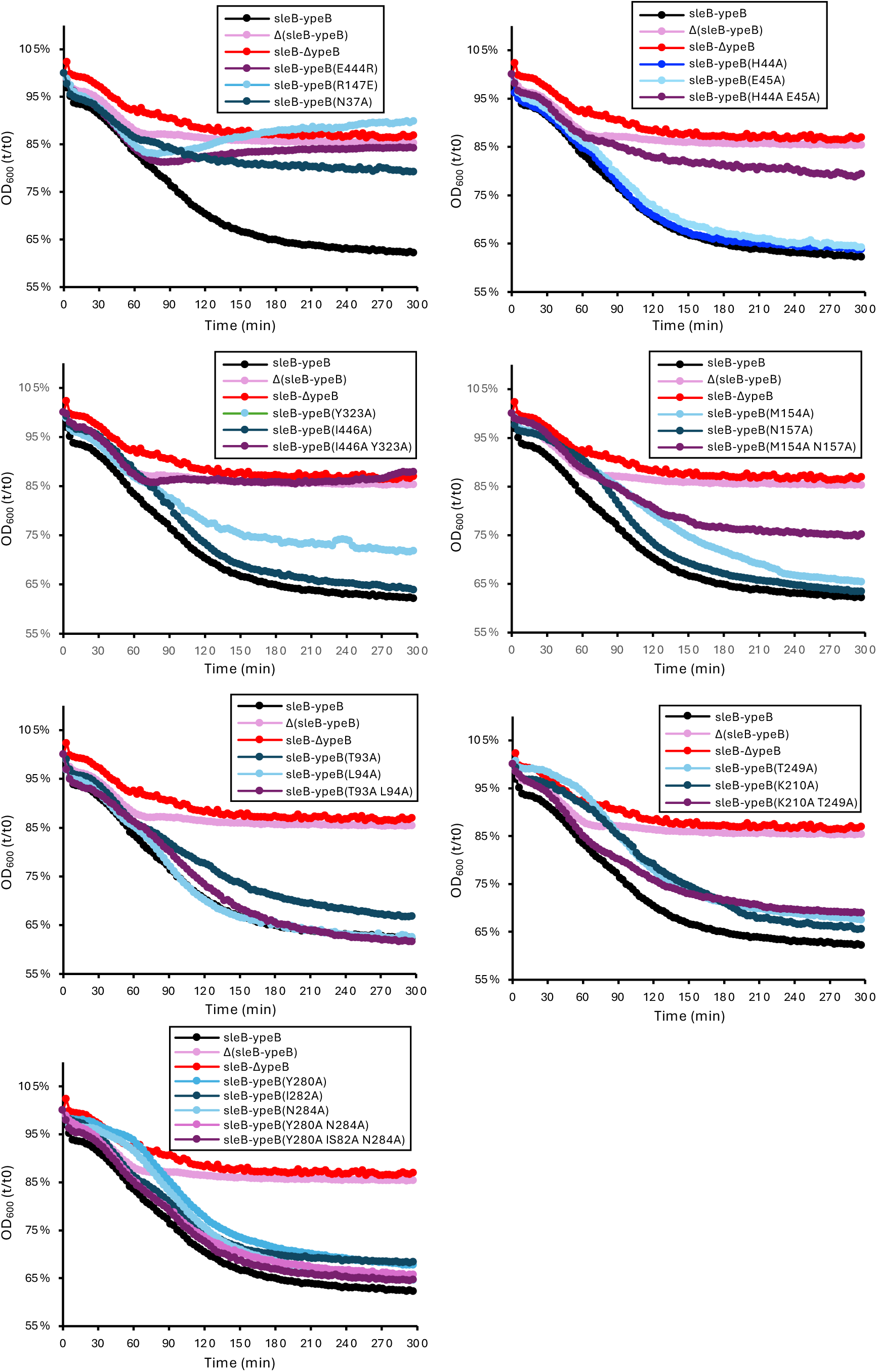
Analysis of spore germination of the YpeB mutants. Representative graphs of spore germination in response to L-alanine as assayed by a reduction in optical density (OD_600_). Histodenz-purified phase-bright spores of the indicated strains lacking *cwlJ*, were induced to germinate with 10 mM L-alanine, and the OD_600_ was monitored over time. The data are plotted as the percent reduction in OD_600_ relative to time 0. In two cases, *ypeB*(M154A, N157A) and *ypeB*(K210A, T249A), the germination defects were more modest than expected based on the sporulation efficiencies in Figure 3B. Spores used for germination were generated on sporulation agar plates at 30°C while the sporulation efficiency was determined from cells sporulated in liquid sporulation medium at 37°C, potentially explaining the discrepancies.

**Figure S6.**
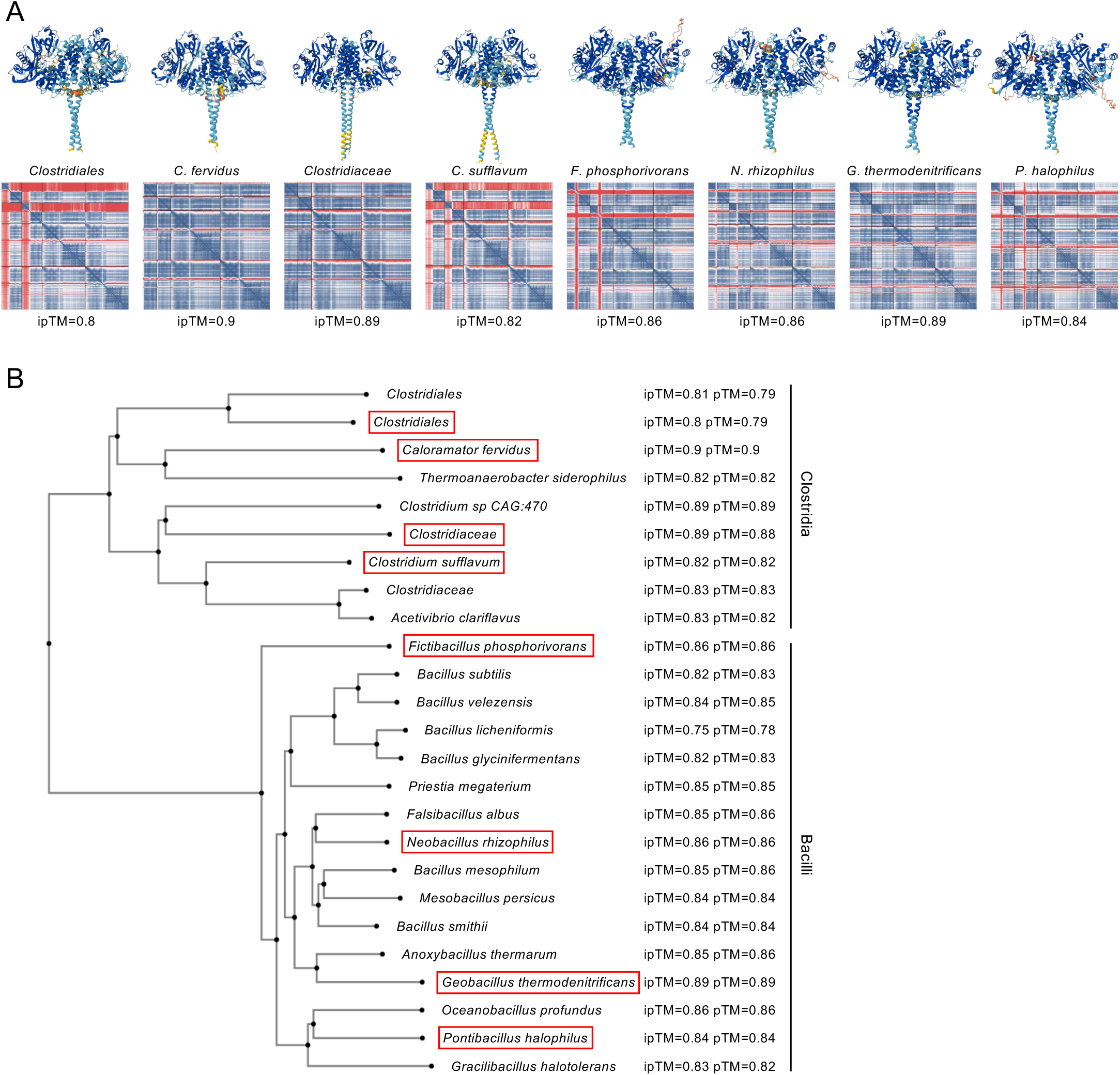
The structure of the *B. subtilis* YpeB_2_-SleB_2_ complex is likely conserved across diverse endospore formers in the Bacillota. (**A**) AlphaFold3-predicted structures of YpeB_2_-SleB_2_ tetrameric complexes from 8 endospore forming bacteria. The models are colored based based on the predicted local distance difference test (pLDDT), a per-residue measure of local confidence. The higher confidence scores are dark blue. Plots of the predicted alignment error (pAE) in Å of all residues against all residues for the top-ranked models are shown below. Low error (blue) corresponds to well-defined relative domain positions. The interface predicted template modeling score (ipTM) for each model is shown below the pAE plot. **(B)** Phylogenetic tree of spore-forming Clostridia and Bacilli for which a YpeB_2_-SleB_2_ tetrameric complex was predicted by AlphaFold3. The ipTM and template modeling (pTM) score for the top model is indicated to the right of the species. Uniprot IDs for all proteins analyzed can be found in Table S4. The predicted structures and pAE plots of the YpeB_2_-SleB_2_ complexes for the species boxed in red are shown in (A).

**Figure S7.**
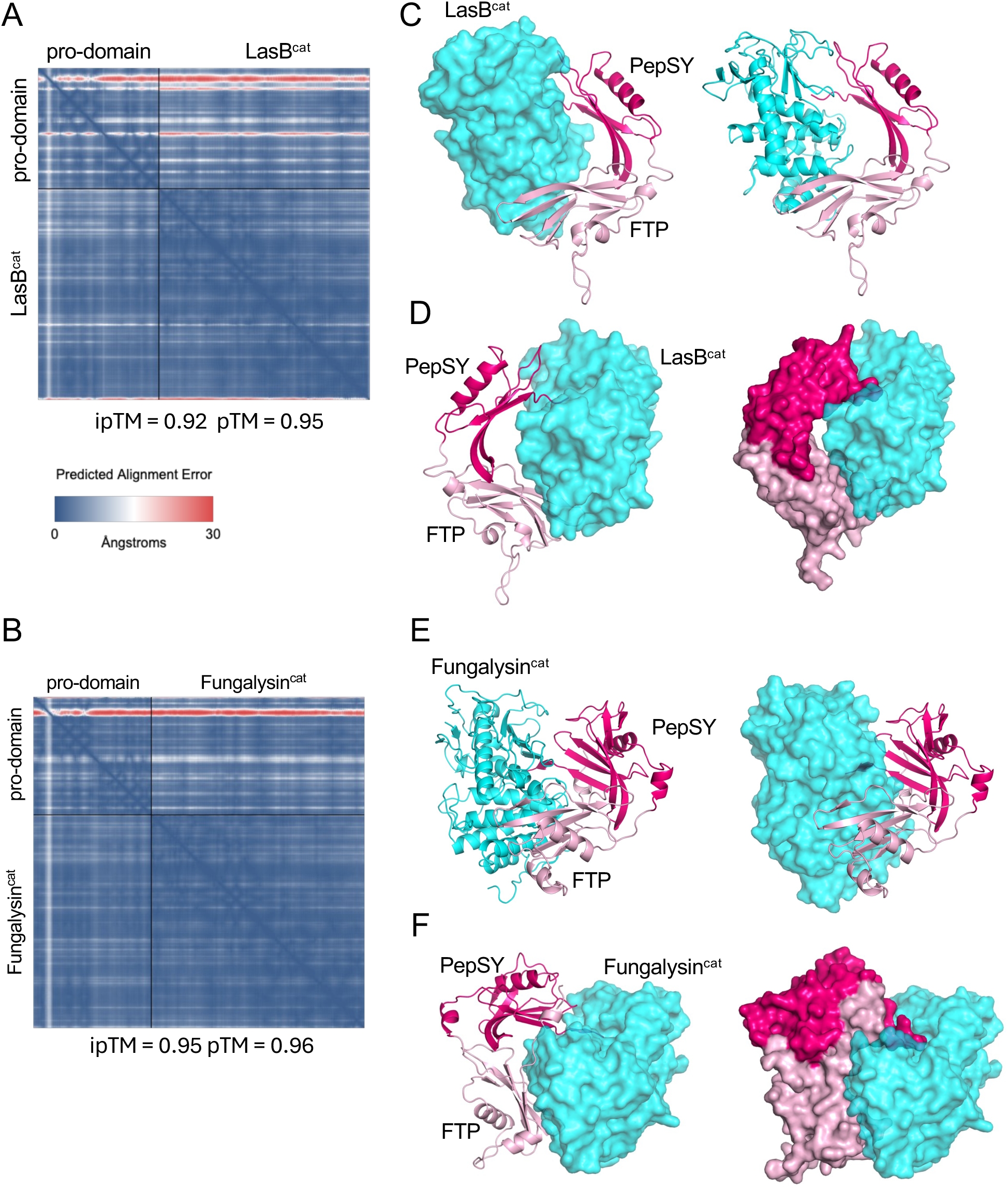
LasB and Fungalysin pro-domains composed of PepSY and FTP domains embrace their catalytic domains. (**A-B**) Plots of the predicted alignment error (pAE) in Å of the AlphaFold3 model of LasB and Fungalysin and their cleaved pro-domains. **(C)** Predicted structure of LasB’s catalytic domain (LasB^cat^) and its cleaved pro-domain. The PepSY domain (dark pink), FTP domain (light pink), and LasB^cat^ (cyan) are rendered as ribbons and surface. **(D)** A rotated view of the predicted complex highlighting the “embrace” of the catalytic domain by both PepSY and FTP domains. **(E)** Predicted structure of Fugalysin’s catalytic domain (Fungalysin^cat^) and its cleaved pro-domain. The PepSY domain (dark pink), FTP domain (light pink), and Fungalysin^cat^ (cyan) are rendered as ribbons and surface. **(D)** A rotated view of the predicted complex highlighting the “embrace” of the catalytic domain by both PepSY and FTP domains.

**Figure S8.**
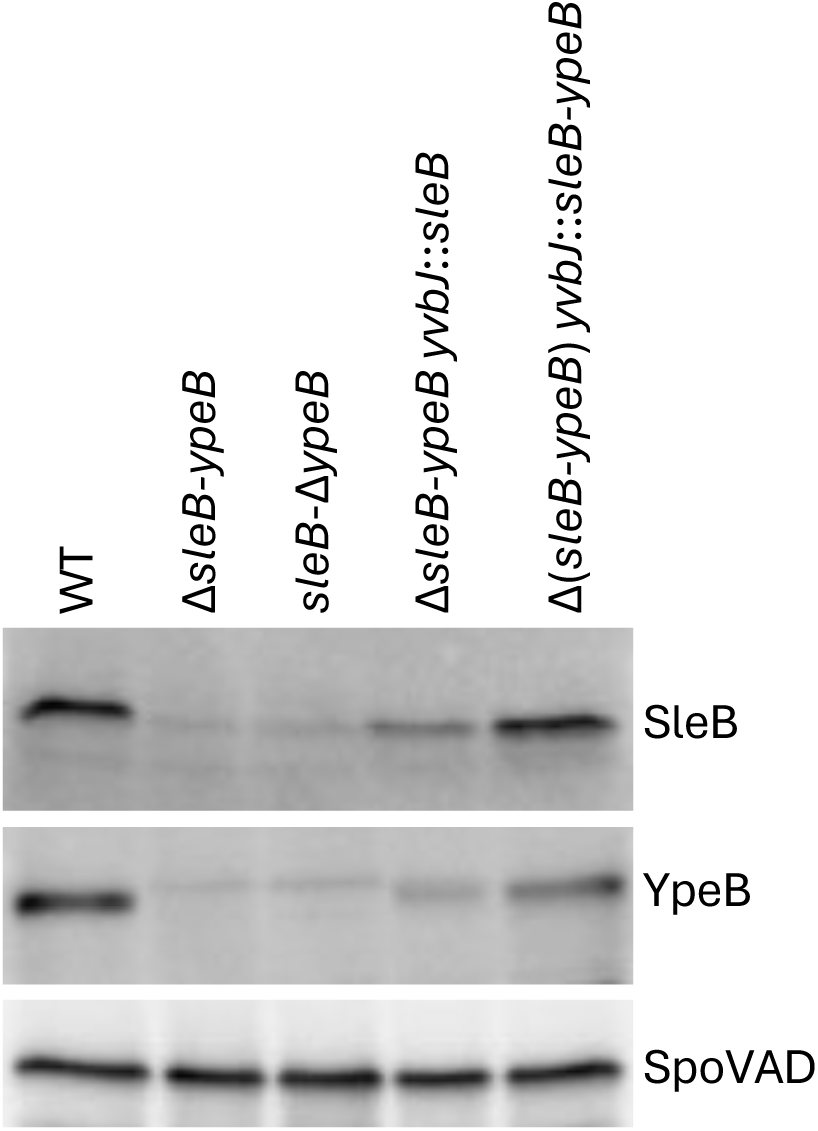
The levels of SleB and YpeB in dormant spores are influenced by the presence of the two genes in a single operon. Representative immunoblots of SleB and YpeB from spore lysates of the indicated strains. SpoVAD controls for loading.

